# Using Shared Features Improves Metabolite Effect Estimation

**DOI:** 10.64898/2026.07.31.742124

**Authors:** Harsh Vardhan Dubey, Gregory Farage, Śaunak Sen

## Abstract

External biological knowledge provides valuable information about relationships among metabolites, yet this information is usually not incorporated directly into statistical estimation procedures. Most existing approaches estimate metabolite effects independently, ignoring known biochemical structure such as shared subclasses and pathway membership. We propose a Bayesian hierarchical framework that improves metabolite effect estimates by incorporating external biological information describing relationships among metabolites. The proposed method improves metabolite-specific estimates by allowing related metabolites to borrow information from one another while preserving metabolite-level inference. We evaluate the methodology using simulation studies across a range of sample sizes and heterogeneity regimes together with three metabolomics applications involving distinct biological annotation structures. Across both simulated and real datasets, incorporating external biological information consistently improves metabolite effect estimation. Gains are most pronounced when sample sizes are small and metabolite classes are informative, i.e. more homogenous within classes.

## 1. Introduction

Metabolomics seeks to comprehensively quantify the small-molecule metabolites present in biological samples, providing a direct biochemical snapshot of cellular physiology. Because metabolites are downstream products of genetic regulation, protein activity, environmental exposure, diet, medication use, and disease processes, metabolomic profiles provide a functional readout of biological state that is often closer to phenotype than genomic or transcriptomic measurements (1, 2). Consequently, metabolomics has become an important component of modern systems biology and precision medicine, enabling biomarker discovery, pathway characterization, and the study of disease-associated metabolic dysregulation.

Rapid advances in mass spectrometry and nuclear magnetic resonance technologies have substantially increased both the sensitivity and throughput of metabolomic experiments. Modern studies routinely quantify hundreds to thousands of metabolites across large collections of biological samples, generating high-dimensional data matrices with complex dependence structures. Large-scale metabolomics workflows now support reproducible profiling of serum, plasma, and other biofluids using liquid chromatography–mass spectrometry and gas chromatography–mass spectrometry platforms (3). These advances have made metabolomics increasingly valuable for population studies, clinical research, and multiomics investigations.

Despite these technological advances, extracting reliable biological insights from high-dimensional metabolomics data remains a major statistical challenge (4, 5). Metabolite abundances are often highly correlated because metabolites participate in shared biochemical pathways, arise from common biosynthetic origins, or belong to related chemical classes. In addition, metabolomics datasets frequently exhibit heterogeneous measurement variability, missing or censored observations, batch effects, and a large number of response variables relative to the number of biological samples. These characteristics make reliable estimation of metabolite-specific associations challenging, particularly when the goal is to estimate effects for many metabolites simultaneously.

Many analyses seek to estimate the association between individual characteristics and each metabolite, and the biological structure linking metabolites is often treated only as down-stream annotation rather than as part of the estimation procedure. This can lead to noisy estimates, especially for metabolites with high measurement variability or limited effective sample size. The availability of curated biochemical classifications and metabolite annotations provides an opportunity to improve inference by incorporating known relationships among metabolites directly into the statistical model. That is the central challenge tackled by our work.

The limitation of estimating metabolite effects independently is not unique to metabolomics. Similar estimation problems arise throughout statistics whenever multiple related parameters must be estimated simultaneously from noisy observations. Bayesian hierarchical models provide a principled framework for addressing this problem by allowing related parameters to share information through a common probabilistic hierarchy. Rather than estimating each parameter in isolation, hierarchical models assume that individual effects arise from a common population distribution, thereby enabling partial pooling of information across related groups. This borrowing of information often reduces estimation variance while preserving meaningful differences among individual parameters (6–9).

The statistical foundations of hierarchical modeling have been extensively studied over the past several decades. Early work by Lindley and Smith (6) established Bayesian estimation for hierarchical linear models, while James and Stein (7) and the subsequent empirical Bayes developments of Efron and Morris (8, 10) demonstrated that jointly estimating related parameters can substantially improve estimation accuracy compared with treating each parameter independently. More recently, hierarchical Bayesian models have become standard tools in biostatistics, epidemiology, genetics, and meta-analysis, where borrowing information across related experimental units or studies improves both estimation precision and predictive performance (9, 11).

The same principle is particularly attractive in metabolomics. Metabolites belonging to the same biochemical subclass often participate in related metabolic pathways, share structural similarities, or arise from common biosynthetic processes. Consequently, one may reasonably expect regression coefficients corresponding to biologically related metabolites to exhibit greater similarity than coefficients associated with unrelated metabolites. Existing metabolomics analyses typically exploit these biological annotations only after statistical estimation has been completed, for example during pathway enrichment or biological interpretation. Incorporating this information directly into the estimation procedure offers the potential to improve statistical efficiency while retaining metabolite-specific inference.

Recent advances have demonstrated the growing potential of Bayesian hierarchical modeling for improving statistical inference in metabolomics and related biomedical applications. Gillies et al. (12) proposed a multilevel Bayesian logistic regression framework that improves metabolite effect size estimation through hierarchical shrinkage while simultaneously accounting for uncertainty arising from missing value imputation. More generally, Busatto and van de Wiel (13) developed a Bayesian horseshoe regression framework that uses external co-data to guide shrinkage of regression coefficients in high-dimensional settings. Molinari et al. (14) introduced a Bayesian dynamic network model for estimating metabolite association networks in cardiovascular disease using hierarchical shrinkage priors, while Collier et al. (15) investigated Bayesian hierarchical meta-regression models for combining evidence across heterogeneous clinical trials. Collectively, these studies illustrate the increasing role of hierarchical Bayesian methodology in improving statistical inference by leveraging shrinkage, auxiliary information, and hierarchical borrowing across diverse biomedical applications.

Despite these advances, the statistical objectives and modeling strategies of these methods differ fundamentally from the problem considered here. Gillies et al. (12) perform global shrinkage of metabolite-specific regression coefficients toward a common prior distribution within Bayesian logistic regression, Busatto and van de Wiel (13) incorporate external co-data into predictor-specific shrinkage priors, Molinari et al. (14) estimate metabolite association networks, and Collier et al. (15) develop hierarchical meta-regression models for combining evidence across studies. In contrast, our objective is to improve metabolite-specific regression coefficient estimates by incorporating external biological information directly into a Bayesian hierarchical model. Rather than shrinking all metabolite effects toward a common global distribution, the proposed framework performs biologically informed partial pooling by allowing metabolites within the same biochemical group to borrow information from one another while preserving metabolite-specific inference. The proposed framework retains conditional independence of metabolite effects given their shared biological subclass while allowing related metabolites to borrow information through a hierarchical prior. To the best of our knowledge, no existing methodology performs Bayesian hierarchical partial pooling of metabolite effect estimates using curated metabolite annotations.

Motivated by this observation, we propose a Bayesian hierarchical method that operates directly on metabolite-specific coefficient estimates and their corresponding standard errors. Metabolites belonging to the same biochemical subclass are assumed to share a common latent distribution, allowing biologically related metabolites to borrow information from one another through partial pooling while preserving metabolite-specific inference. Posterior inference is performed using an efficient Gibbs sampler based on the auxiliary-variable representation of the Half-Cauchy prior proposed by Makalic and Schmidt (16).

The primary contributions of this paper are:

1. We develop a statistical framework for incorporating external biological information into metabolite effect estimation through Bayesian hierarchical modeling.
2. Through extensive simulation studies, we demonstrate consistent improvements in coefficient estimation accuracy and predictive performance across a range of sample sizes and heterogeneity regimes.
3. We evaluate the proposed methodology on three metabolomics data sets, demonstrating improved estimate reproducibility and predictive performance across studies with substantially different biological annotation structures.
4. We develop an efficient Bayesian inference procedure based on a fully conjugate Gibbs sampler using the Makalic and Schmidt (16) auxiliary-variable representation of the Half-Cauchy prior.
5. More broadly, we illustrate a general framework for incorporating external biological knowledge into matrix-based regression models, providing a foundation for future extensions based on richer biological ontologies and learned molecular representations.

The remainder of the paper is organized as follows. Section 2 introduces the proposed Bayesian hierarchical framework and describes the posterior inference procedure. Section 3 presents the simulation design together with the three metabolomics datasets, their preprocessing procedures, and the external biological information incorporated into the hierarchical model. Section 4.1 presents the simulation results, while Sections 4.2–4.4 evaluate the proposed methodology using the three metabolomics applications. Finally, Section 5 concludes with a discussion of the findings, limitations, and directions for future research.

## 2. Methods

### 2.1. Motivation for Hierarchical Borrowing of Information

Many statistical procedures produce an estimated effect and corresponding standard error for each metabolite separately. Consequently, inference for metabolite *j* depends only on the observations associated with that metabolite, and no information is shared across metabolites during estimation. This independence simplifies estimation and preserves metabolite-specific interpretation, but it does not exploit external biological knowledge describing relationships among metabolites.

Extensive external biological information describing relationships among metabolites is already available through curated resources such as the Human Metabolome Database (HMDB) and the Kyoto Encyclopedia of Genes and Genomes (KEGG) (17, 18), where metabolites are organized into superclasses, subclasses, lipid families, and biochemical pathways. These annotations represent valuable prior biological knowledge that is typically used only after statistical analysis for pathway enrichment and biological interpretation, rather than during estimation itself. The central objective of the proposed methodology is to incorporate this external biological information directly into metabolite effect estimation. Specifically, we introduce a Bayesian hierarchical model in which metabolites belonging to the same biological subclass share a common latent subclass effect, allowing biologically related metabolites to borrow information through partial pooling. Importantly, metabolite-specific effects remain conditionally independent given the shared subclass-specific latent effect, thereby preserving metabolite-level inference while improving estimation stability and, consequently, downstream prediction.

To improve estimation, we consider a framework that allows related metabolites to share information. Specifically, suppose that each metabolite belongs to one of *H* biological groups. Let

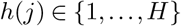

denote the group membership of metabolite *j*. Rather than viewing metabolite effects as completely unrelated, we assume that metabolites within the same group arise from a common distribution centered around a group-specific mean effect.

Under this perspective, the observed effect estimate 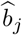 is regarded as a noisy measurement of an underlying true effect *θ*_*j*_. Metabolites belonging to the same biological group are assumed to have related true effects, allowing information to be shared across metabolites within that group. Group-level effects are then connected through a global population mean, producing a hierarchical structure that enables borrowing of information at multiple levels.

The resulting framework provides several advantages. First, effect estimates for metabolites with large standard errors are stabilized through shrinkage toward the corresponding group mean. Second, group-level estimates are themselves informed by all metabolites assigned to the group, improving estimation when the number of metabolites per group is moderate or large. Finally, the hierarchical structure naturally accounts for uncertainty at each level of the model, yielding posterior distributions that reflect both within-group and between-group variability.

This hierarchical borrowing mechanism is conceptually similar to empirical Bayes and random-effects models, where information is shared across related units to improve estimation accuracy (19). In the present setting, the hierarchy exploits known biological groupings of metabolites to obtain more stable and interpretable estimates of covariate effects than can be achieved by analyzing each metabolite independently.

The Bayesian hierarchical model used to accomplish this borrowing of information is described in Section 2.2.

### 2.2. Bayesian Hierarchical Model

For a fixed covariate of interest, let 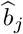 denote the effect estimate for metabolite *j*, and let *s*_*j*_ denote its corresponding estimated standard error. We assume that the input effect estimates are unbiased for their underlying metabolite-specific effects and are uncorrelated across metabolites, as is the case for the metabolite-specific coefficient estimates obtained from a Matrix Linear Model. Accordingly, each 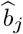 is treated as an observed summary statistic representing a noisy measurement of an unobserved true metabolite effect *θ*_*j*_, with sampling variability characterized by 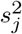.

Suppose that metabolite *j* belongs to biological group *h*(*j*), where

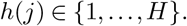

The first level of the hierarchy models the observed estimate as

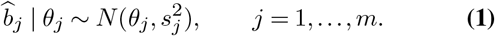

Equation (1) treats the standard error *s*_*j*_ as known and accounts for the uncertainty associated with estimation of the metabolite-specific effect. Conditional on the true effect *θ*_*j*_, the observed estimate 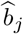 is assumed to follow a normal distribution centered at *θ*_*j*_.

To allow borrowing of information among biologically related metabolites, the latent effects are assumed to arise from a group-specific distribution,

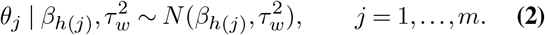

Here, *β*_*h*_ represents the mean effect associated with biological group *h*, while 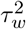 quantifies the variability of metabolite effects within a group. Small values of 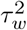 imply that metabolites within the same group exhibit similar effects and therefore experience stronger shrinkage toward the corresponding group mean. The proposed hierarchy induces dependence among metabolite effect estimates only through the shared subclass-specific latent parameter *β*_*h*_. Conditional on *β*_*h*_, the metabolite-specific effects *θ*_*j*_ are mutually independent. Consequently, the Bayesian hierarchy preserves the interpretation of individual metabolite effects while allowing external biological information to stabilize estimation through partial pooling.

At the next level of the hierarchy, group-specific effects are modeled as

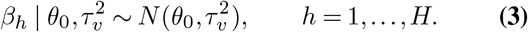

The parameter *θ*_0_ represents an overall population-level effect shared across all metabolite groups, while 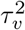 governs the variability between groups. Smaller values of 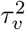 induce stronger shrinkage of group means toward the global mean. The complete hierarchical structure may therefore be summarized as

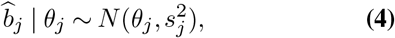

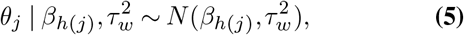

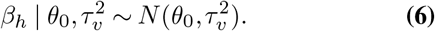

The conditional relationships among the observed estimates, metabolite-specific effects, subclass-level effects, and overall mean are summarized in Fig. 1.

**Fig. 1.**
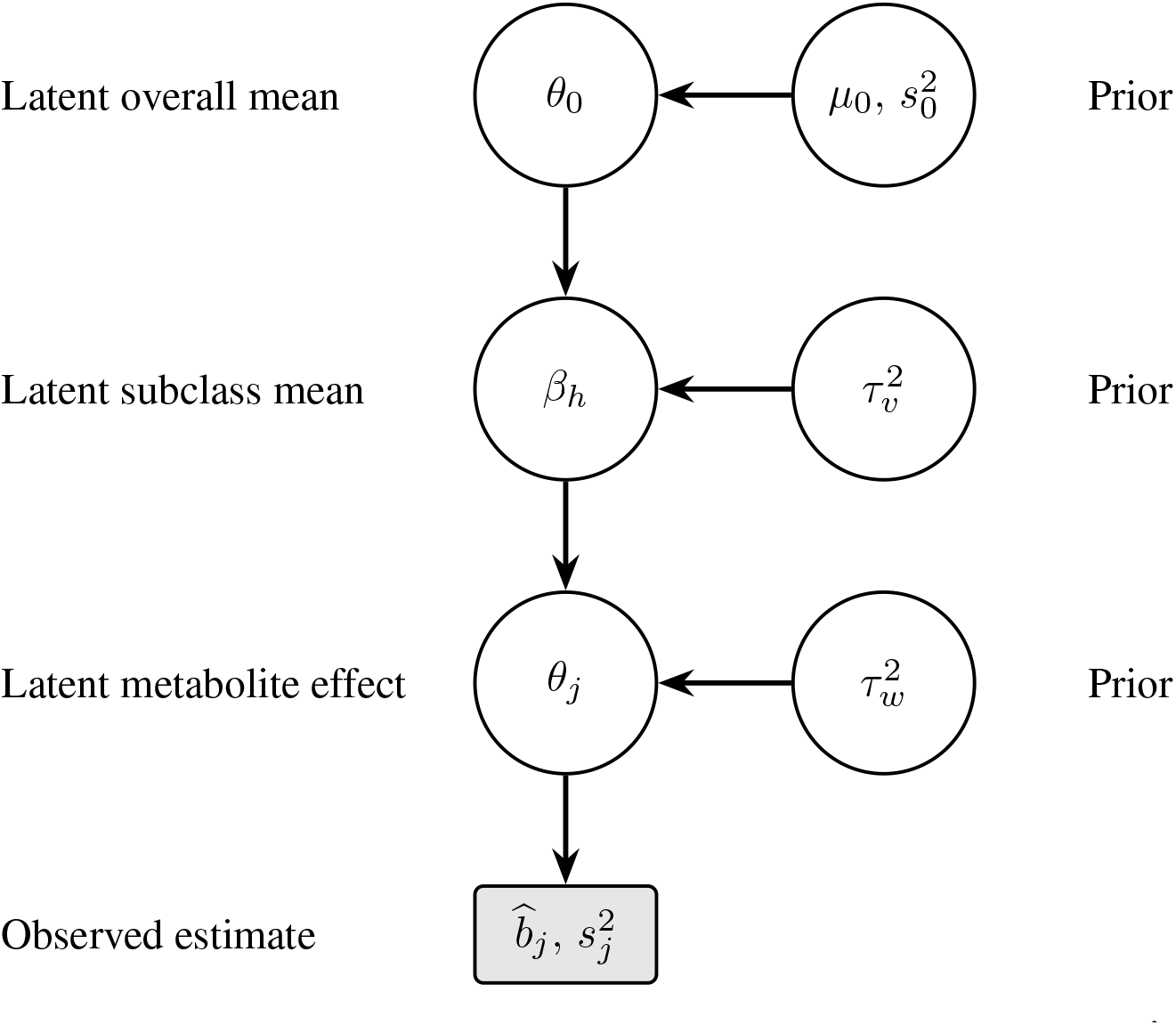
Graphical representation of the proposed Bayesian hierarchical model for a fixed covariate. The observed metabolite estimate 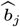 and its estimated variance 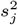 provide information about the latent metabolite-specific effect *θ*_*j*_. Metabolites belonging to subclass *h* share the latent subclass effect *β*_*h*_, which permits biologically informed partial pooling, while the subclass effects are centered around the overall mean *θ*_0_. The variance parameters 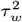 and 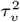 control within-subclass and between-subclass heterogeneity, respectively. Metabolite-specific effects are conditionally independent given their subclass-specific latent effects.

The proposed hierarchy serves as a statistical mechanism for incorporating external biological information into the estimation procedure rather than treating metabolite effects as independent quantities. This hierarchical formulation induces partial pooling across metabolites belonging to the same biological group while simultaneously allowing information to be shared across groups through the global mean parameter *θ*_0_. Consequently, estimates for metabolites with limited information are stabilized by borrowing strength from related metabolites, while metabolites with strong individual signals remain largely determined by their own data.

The proposed hierarchy defines a Bayesian model that incorporates external biological information through biologically informed partial pooling while preserving metabolite-specific inference. Prior distributions, posterior inference, and the Gibbs sampling algorithm used for posterior computation are provided in Appendices A.1–A.4. We next describe how posterior samples are summarized to obtain metabolite effect estimates.

### 2.3. Posterior Effect Estimates

The primary quantity of interest in the proposed hierarchical model is the metabolite-specific effect parameter *θ*_*j*_, which represents the underlying effect of the covariate of interest on metabolite *j* after borrowing information from biologically related metabolites. Following Gibbs sampling, posterior inference for *θ*_*j*_ is based on the collection of retained Markov chain samples

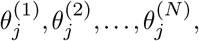

obtained after discarding burn-in iterations. The posterior mean is used as the primary point estimator and is given by

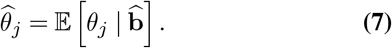

Because the posterior expectation is generally unavailable in closed form, it is approximated using the Monte Carlo average of the retained posterior samples (20),

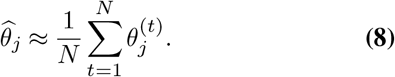

The estimator in Equation (8) serves as the final Bayesian effect estimate reported throughout the remainder of the paper. In addition to point estimation, posterior uncertainty may be quantified using the empirical posterior distribution of the sampled values. For example, a 100(1 − *α*)% credible interval may be obtained from the corresponding posterior quantiles,

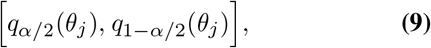

where *q*_*p*_(*θ*_*j*_) denotes the *p*th posterior quantile.

The posterior mean estimator combines information from three sources: the observed metabolite estimate 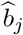, the effects of other metabolites within the same biological group, and the global population-level effect. Consequently, the resulting estimates exhibit adaptive shrinkage toward biologically informed group means while preserving strong metabolite-specific signals when supported by the data.

Throughout this paper, all reported Bayesian metabolite effect estimates correspond to the posterior means defined in Equation (8).

## 3. Simulation Design and Datasets

The proposed methodology was evaluated using both simulated data and three metabolomics applications representing distinct biological settings. This section describes the simulation design, the real-world datasets, the associated preprocessing procedures, and the external biological information incorporated into the Bayesian hierarchical model.

### 3.1. Simulation Design

We conducted a simulation study to evaluate the benefit of incorporating external biological information through the proposed hierarchical model using coefficient estimates and corresponding standard errors obtained from the `MatrixLM` Julia package used by Farage et al. (21). The `MatrixLM` package serves as the baseline method in all simulation and real-data comparisons. More generally, the proposed Bayesian hierarchical framework is independent of any particular upstream estimation procedure and can be applied to metabolite effect estimates and their associated standard errors obtained from a wide range of statistical models.

Data were generated from a Linear Model,

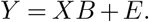

where *Y* ∈ ℝ^*n*×*m*^ is the simulated metabolite response matrix, *X* ∈ ℝ^*n*×*p*^ is the subject-level covariate matrix, *B* ∈ ℝ^*p*×*m*^ is the true coefficient matrix, and *E* ∈ ℝ^*n*×*m*^ is the random error matrix.

In all simulations, the entries of *X* were generated independently from a standard normal distribution. The entries of the error matrix were generated independently as

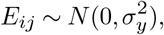

with *σ*_*y*_ = 1.

The true coefficient matrix *B* was generated using a one-level hierarchical structure over metabolite subclasses. Specifically, for each covariate *k* = 1, …, *p*, a global covariate-specific effect was generated as

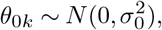

where *σ*_0_ = 0.2. For each metabolite subclass *h* = 1, …, *H*, the subclass-level effect for covariate *k* was generated as

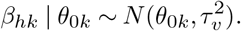

Finally, for metabolite *j* belonging to subclass *h*(*j*), the true metabolite-specific coefficient was generated as

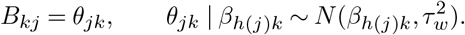

Thus, *τ*_*v*_ controls between-subclass heterogeneity, while *τ*_*w*_ controls within-subclass heterogeneity. Larger values of *τ*_*v*_ imply that subclass-level mean effects may differ substantially from one another. Larger values of *τ*_*w*_ imply greater variability among individual metabolite effects within the same subclass.

We considered *p* = 10 covariates and *H* = 4 metabolite sub-classes. The subclass proportions were fixed at

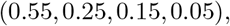

so that the simulated metabolite classes were intentionally unbalanced. This reflects the fact that, in real metabolomics datasets, biological or chemical subclasses often contain unequal numbers of metabolites.

Three levels of hierarchical heterogeneity were considered to evaluate the robustness of the proposed Bayesian framework across varying degrees of metabolite-effect variability. The overall hierarchical heterogeneity is governed by

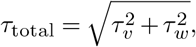

which represents the marginal standard deviation of metabolite effects around the global mean under the hierarchical model.

The between-subclass and within-subclass standard deviations were specified as

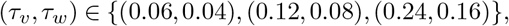

corresponding to low (*τ*_*total*_ = 0.072), moderate (*τ*_*total*_ = 0.144), and high (*τ*_*total*_ = 0.288) heterogeneity respectively. These values were obtained by proportionally scaling both variance components while maintaining a constant ratio

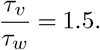

Consequently, the proportion of total metabolite-effect variability attributable to between-subclass and within-subclass variation remained unchanged across the three regimes, while the overall magnitude of hierarchical heterogeneity increased. This design isolates the effect of increasing heterogeneity without simultaneously altering the underlying hierarchical structure or the relative similarity of metabolites within the same biochemical subclass.

To investigate the effect of sample size on estimation and prediction performance, the number of metabolites was fixed at

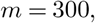

while the sample size varied over

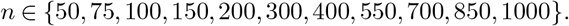

The choice of *m* = 300 was motivated by preliminary simulation studies, which indicated that the relative performance of the proposed Bayesian framework was largely insensitive to the number of metabolites. Fixing the number of metabolites therefore allowed the simulation study to isolate the effect of sample size. These settings span the range from small metabolomics studies, where hierarchical borrowing is expected to provide the greatest benefit, to large studies in which metabolite effects can be estimated with substantially greater precision from the observed data alone.

For each combination of heterogeneity regime and sample size, we generated 100 independent simulated datasets. Each dataset was randomly divided into training and testing sets using a 70/30 split across subjects. `MatrixLM` was fit on the training data to obtain coefficient estimates 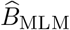 and the corresponding standard error estimates. The proposed Bayesian hierarchical model was then applied separately to the estimated effects for each covariate to obtain posterior mean estimates 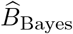.

Coefficient estimation accuracy was evaluated using the mean squared error between the estimated and true coefficient matrices,

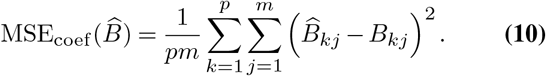

Prediction accuracy was evaluated on the held-out testing data. If *X*_test_ and *Y*_test_ denote the test-set covariate and response matrices, the prediction mean squared error was defined as

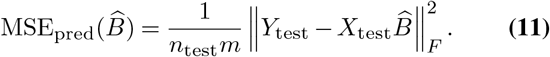

For both coefficient estimation and prediction, performance was summarized using the MSE ratio

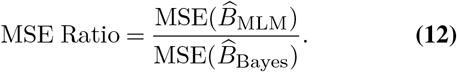

Values greater than one indicate improved performance of the proposed Bayesian hierarchical estimator relative to `MatrixLM`. Ratios were averaged over the 100 independent simulation replicates within each scenario.

### 3.2. COPDGene Study

The first real-data application considered the COPDGene metabolomics study originally analyzed by Gillenwater et al. (22) and subsequently used to illustrate the matrix linear model framework by Farage et al. (21). The original study analyzed the COPDGene and SPIROMICS cohorts to investigate sex-specific metabolomic differences associated with chronic obstructive pulmonary disease (COPD), with the COPDGene cohort consisting of 839 participants and 999 measured metabolites. Plasma samples were profiled using the Metabolon Global Metabolomics Platform, which provides broad coverage of the human metabolome and is widely used for biomarker discovery because of the accessibility and minimally invasive nature of plasma sampling (22). The extracted samples were partitioned into four distinct fractions for comprehensive analysis. Two fractions were allocated for analysis using reverse-phase/ultrahigh performance liquid chromatography-tandem mass spectrometry (RP/UPLC-MS/MS) equipped with positive ion mode electrospray ionization (ESI). Another fraction was dedicated to RP/UPLC-MS/MS analysis but utilized negative ion mode ESI. The fourth fraction was reserved for analysis through hydrophilic interaction chromatography (HILIC)/UPLC-MS/MS, again employing negative ion mode ESI.

COPD case status was defined using established spirometric criteria corresponding to at least moderate airflow obstruction, namely a post-bronchodilator forced expiratory volume in one second to forced vital capacity ratio (FEV_1_/FVC) below 0.50 together with a forced expiratory volume in one second percent predicted (FEV_1_pp) below 80%. Control subjects were defined by a FEV_1_/FVC ratio greater than 0.70 and a FEV_1_pp exceeding 80% (22). Characterizing metabolomic differences between COPD cases and controls is important for improving our understanding of disease mechanisms and for identifying potential biomarkers relevant to precision medicine.

Following the preprocessing pipeline described by Farage et al. (21), missing metabolite measurements were imputed using *k*-nearest neighbor imputation, probabilistic quotient normalization was applied to reduce dilution effects (23), and metabolite abundances were subsequently log_2_ transformed to improve symmetry and stabilize the variance. After pre-processing, the final dataset consisted of 784 individuals and 999 quantified metabolites. Each metabolite was annotated using one of 106 biochemical subclasses obtained from the Metabolon metabolite annotation database. These sub-class labels served as the biological grouping variable in the proposed Bayesian hierarchical model, allowing metabolite-specific effects to borrow information from chemically related metabolites within the same subclass. The metabolite subclasses therefore provide biologically meaningful external information that can be incorporated into the proposed hierarchical model for effect estimation.

To investigate how the benefit of biologically informed partial pooling depends on the amount of available data, we conducted a repeated subsampling analysis of the COPDGene study. Training subsets of varying sizes,

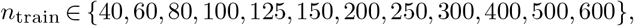

were repeatedly sampled without replacement, with the remaining subjects serving as an independent test set. For each training subset, `MatrixLM` was fit to obtain metabolite-specific coefficient estimates and their corresponding standard errors for each covariate; the coefficient estimates are denoted by 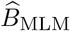. The proposed Bayesian hierarchical model was subsequently applied to these estimates to obtain posterior mean estimates, denoted by 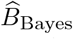. Posterior inference was performed using a Gibbs sampler with 5,000 iterations, of which the first 1,000 were discarded as burn-in. The prior hyperparameters of the hierarchical model were set to *µ*_0_ = 0 and *s*_0_ = 1.

Because the true metabolite effects are unknown in real data, direct estimation error cannot be evaluated. Instead, the `MatrixLM` estimates obtained from the independent test subset were used as a common reference against which both the training `MatrixLM` estimates and the Bayesian posterior mean estimates were compared. For each covariate, we computed the corresponding mean squared errors,

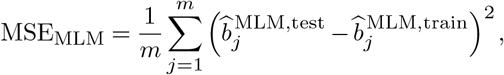

and

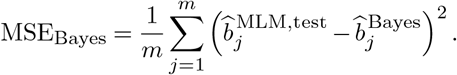

These quantities were summarized using the *Reference MSE Ratio* (RMR),

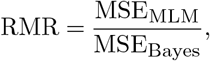

which measures the relative agreement of the two estimators with the independent reference estimates. Values of RMR *>* 1 indicate that the Bayesian hierarchical estimator produces metabolite effect estimates that are closer to the independent reference estimates than those obtained directly from `MatrixLM`, thereby demonstrating improved estimation stability through biologically informed partial pooling.

### 3.3. Statin-Associated Muscle Symptoms (SAMS) Study

The second real-data application considered the Statin-Associated Muscle Symptoms (SAMS) study originally analyzed by Garrett et al. (24) and subsequently included as one of the motivating datasets for the `MatrixLM` package by Farage et al. (21). While the original study focused on metabolomic and lipidomic changes associated with statin rechallenge, Farage et al. (21) considered a complementary scientific question by investigating the association between fish oil supplementation and triglyceride composition. This analysis illustrates how matrix linear models can be used to study treatment-related effects beyond the primary objective of the original clinical investigation.

Plasma lipidomic profiles were generated using liquid chromatography–mass spectrometry (LC–MS), resulting in measurements for 770 identified triglycerides from 98 individuals (24). The objective of the analysis was to quantify the association between fish oil supplementation and triglyceride abundances across the lipidome. Following the preprocessing strategy described in Farage et al. (21), missing values were imputed using Quantile Regression Imputation of Left-Censored data implemented in the `imputeLCMD` package (25), probabilistic quotient normalization was applied (23), metabolite abundances were log-transformed to improve distributional symmetry, and batch effects were removed using the ComBat method implemented in the `sva` package (26, 27).

After preprocessing, the final dataset consisted of 98 individuals and 770 triglycerides. Rather than using detailed biochemical subclasses, triglycerides were grouped according to their degree of unsaturation, measured by the number of carbon double bonds. Specifically, four biologically meaningful groups were considered: (i) fewer than three double bonds, (ii) three to five double bonds, (iii) six to eight double bonds, and (iv) at least nine double bonds. These groups provide biologically interpretable external information reflecting structural similarities among triglycerides and were used as the grouping variable in the proposed Bayesian hierarchical model. This application demonstrates that the proposed framework is not restricted to curated metabolite subclass annotations but can also leverage biologically meaningful groupings derived from domain-specific knowledge. As in the COPDGene study, we conducted a repeated sub-sampling analysis to evaluate the stability of metabolite effect estimation under varying amounts of available training data. Owing to the smaller sample size of the SAMS cohort (*n* = 98), training subsets of size

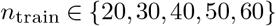

were repeatedly sampled without replacement, with the remaining subjects serving as an independent test set. For each training subset, `MatrixLM` was used to obtain metabolite-specific coefficient estimates and corresponding standard errors, after which the proposed Bayesian hierarchical model was applied to obtain posterior mean estimates. Posterior inference was performed using a Gibbs sampler with 5,000 iterations, of which the first 1,000 were discarded as burn-in. The prior hyperparameters of the hierarchical model were set to *µ*_0_ = 0 and *s*_0_ = 1. Estimation performance was evaluated using the same Reference MSE Ratio (RMR) introduced for the COPDGene analysis, allowing a direct comparison of the relative stability of the two estimators across varying training sample sizes.

### 3.4. PANSteatitis Mozambique Tilapia Study

The third real-data application considered the PANSteatitis Mozambique tilapia study originally conducted by Koelmel et al. (28) and subsequently analyzed using matrix linear models by Farage et al. (21). The original study investigated plasma lipidomic profiles of Mozambique tilapia (*Oreochromis mossambicus*) collected from the Loskop Dam in South Africa to identify lipid biomarkers associated with pansteatitis, a debilitating inflammatory disease of adipose tissue that contributed to large-scale mortality events affecting both fish and Nile crocodiles (*Crocodylus niloticus*) within the region (28). While the primary objective of the original study was to identify lipid markers distinguishing healthy and diseased fish, Farage et al. (21) considered the complementary problem of investigating how fish age influences the composition of different lipid groups.

Plasma lipidomic profiles were generated using liquid chromatography coupled with high-resolution tandem mass spectrometry, resulting in measurements for 590 metabolites from 51 Mozambique tilapia (28). Clinical lipidomics has proven valuable for identifying biomarkers and studying disease mechanisms, but wildlife lipidomics presents additional analytical challenges because metabolite abundances naturally vary with environmental conditions, geography, and age. Consequently, robust statistical methods are particularly important for distinguishing biologically meaningful variation from natural environmental variability.

Following the preprocessing pipeline described by Farage et al. (21), the metabolomic dataset obtained from the Metabolomics Workbench (29) contained no missing observations and therefore required no imputation. Metabolite abundances were normalized using probabilistic quotient normalization (23) and subsequently log_2_ transformed to improve distributional symmetry and stabilize the variance.

After preprocessing, the final dataset consisted of 44 individuals and 590 metabolites. Each metabolite was assigned to one of 13 biochemical subclasses obtained from the available metabolite annotations. These subclass annotations provide biologically meaningful external information describing chemical relationships among metabolites and were used as the grouping variable in the proposed Bayesian hierarchical model. This application illustrates how curated biochemical annotations can be incorporated directly into metabolite effect estimation to improve estimation stability while preserving metabolite-specific inference.

As in the COPDGene and SAMS studies, we conducted a repeated subsampling analysis to evaluate the stability of metabolite effect estimation under varying amounts of available training data. Owing to the smaller sample size of the PANSteatitis study (*n* = 44), training subsets of size

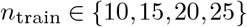

were repeatedly sampled without replacement, with the remaining subjects serving as an independent test set. For each training subset, `MatrixLM` was used to obtain metabolite-specific coefficient estimates and corresponding standard errors, after which the proposed Bayesian hierarchical model was applied to obtain posterior mean estimates. Posterior inference was performed using a Gibbs sampler with 5,000 iterations, of which the first 1,000 were discarded as burn-in. The prior hyperparameters of the hierarchical model were set to *µ*_0_ = 0 and *s*_0_ = 1. Estimation performance was evaluated using the same Reference MSE Ratio (RMR) introduced for the COPDGene analysis, facilitating a consistent comparison of estimation stability across all three metabolomics studies.

## 4. Results

We next evaluate the proposed methodology on the datasets described in Section 3. Performance is assessed from two complementary perspectives. First, we examine the accuracy and reproducibility of metabolite effect estimation. Second, we investigate whether improved estimation translates into improved predictive performance.

### 4.1. Simulation Results

We first evaluate the proposed Bayesian hierarchical framework under controlled simulation settings, where the true regression coefficient matrix is known. This allows the estimation error of both `MatrixLM` and our proposed Bayesian estimator to be assessed directly while simultaneously evaluating predictive performance on independent testing data.

Figure 2 summarizes the coefficient estimation performance across the three heterogeneity regimes. The proposed Bayesian hierarchical estimator consistently outperformed `MatrixLM`, with Estimation MSE Ratios greater than one across all sample sizes and heterogeneity levels. The largest improvements were observed when the available sample size was limited, demonstrating that biologically informed partial pooling is most beneficial when individual metabolite effects are estimated with substantial uncertainty.

**Fig. 2.**
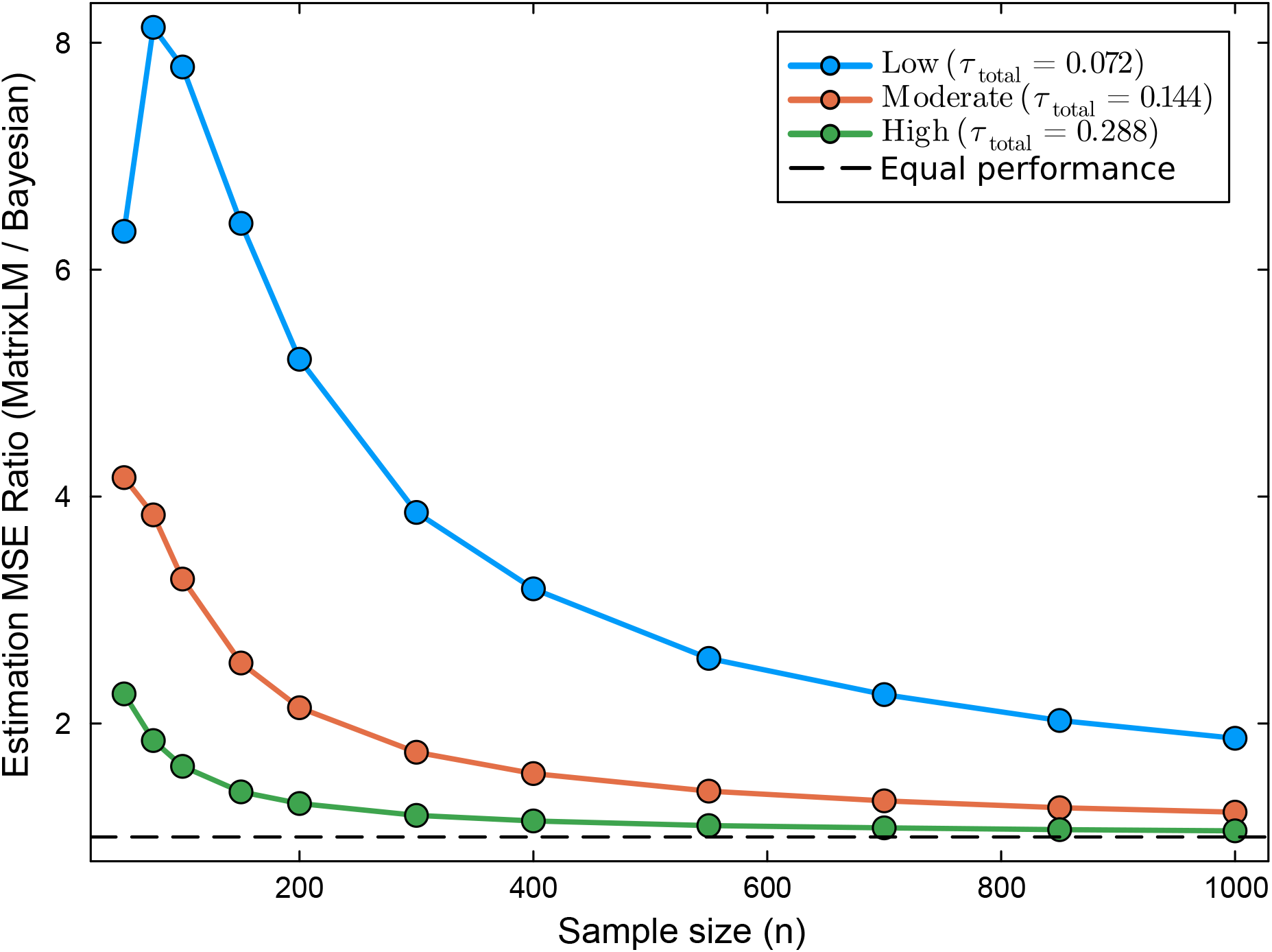
Estimation MSE Ratio as a function of sample size under three levels of hierarchical heterogeneity. The ratio compares the coefficient estimation mean squared errors of `MatrixLM` and the proposed Bayesian hierarchical estimator relative to the true regression coefficients. Across all heterogeneity regimes, the Bayesian estimator consistently achieves lower estimation error than `MatrixLM`, with the greatest improvements observed at smaller sample sizes. The advantage gradually decreases as sample size increases, reflecting the increasing precision of unpooled estimates when more observations are available.

The magnitude of improvement depended on the overall hierarchical heterogeneity. Under the low heterogeneity regime, where metabolites within the same subclass exhibit the greatest similarity, the Bayesian estimator substantially reduced coefficient estimation error, achieving Estimation MSE Ratios exceeding eight for the smallest sample sizes. As the overall heterogeneity increased, the benefit of partial pooling gradually diminished because metabolite effects within the same subclass became increasingly variable. Nevertheless, even under the high heterogeneity regime, the Bayesian estimator consistently produced more accurate coefficient estimates than `MatrixLM`.

Across all three heterogeneity regimes, the Estimation MSE Ratio decreased smoothly toward one as the sample size increased. This behavior is expected because the uncertainty of metabolite-specific estimates decreases with increasing sample size, reducing the additional benefit obtained from borrowing information across biologically related metabolites. Prediction performance is summarized in Figure 3. The proposed Bayesian estimator consistently achieved lower prediction error than `MatrixLM` across all simulation settings, although the improvements were considerably smaller than those observed for coefficient estimation. Prediction MSE Ratios remained greater than one across all sample sizes and heterogeneity regimes, with the largest gains again occurring when the available training sample was smallest.

**Fig. 3.**
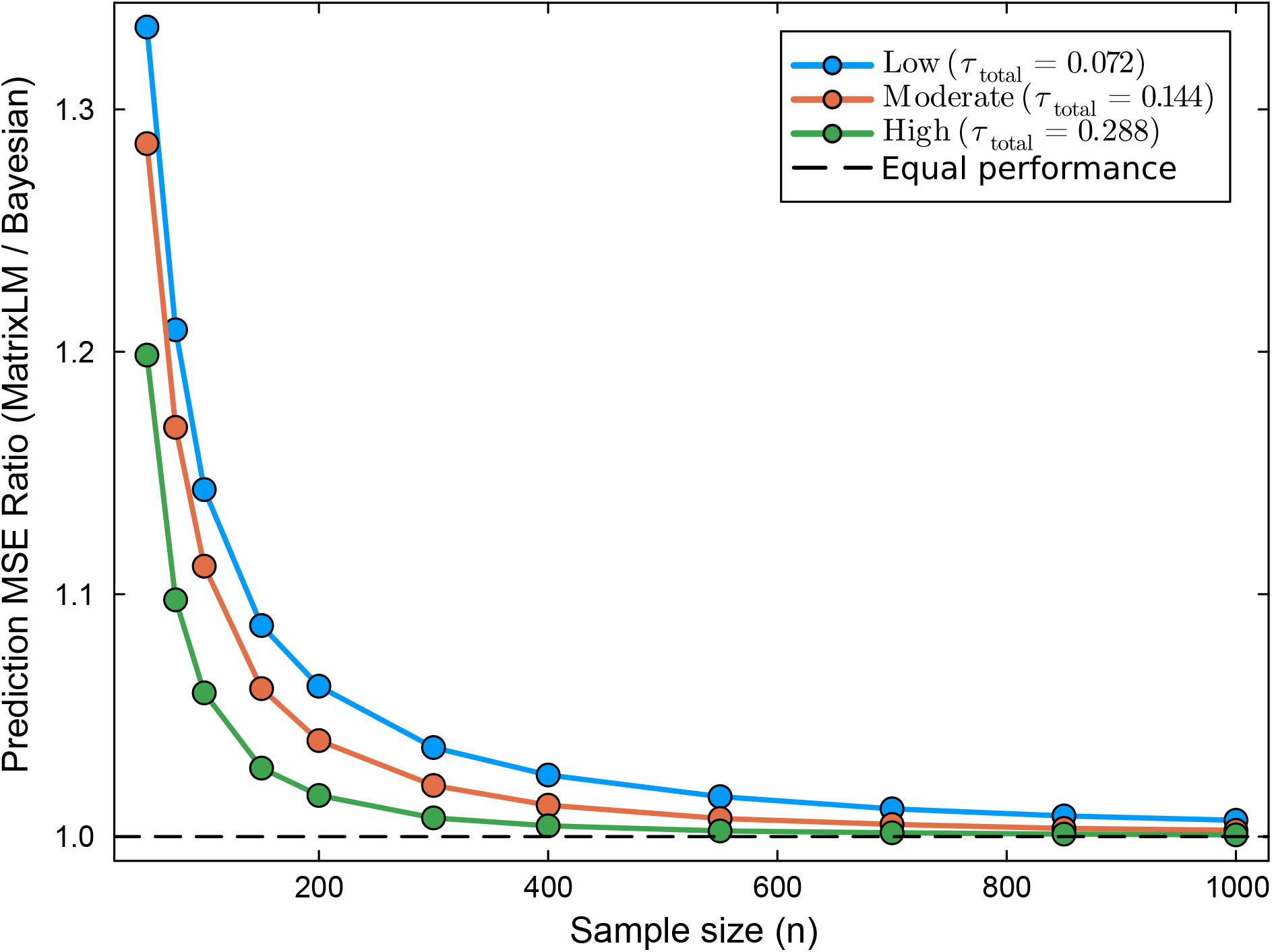
Prediction MSE Ratio as a function of sample size under three levels of hierarchical heterogeneity. The ratio compares the prediction mean squared errors of `MatrixLM` and the proposed Bayesian hierarchical estimator on independent testing data. Although the Bayesian estimator consistently achieves lower prediction error across all heterogeneity regimes, the improvements are more modest than those observed for coefficient estimation and diminish as sample size increases.

The more modest improvements in prediction are expected because prediction depends on the combined contribution of all regression coefficients through the matrix product *XB*. Consequently, substantial improvements in individual coefficient estimates do not necessarily translate into equally large reductions in prediction error. Taken together, these results demonstrate that the primary benefit of incorporating external biological information lies in improving metabolite effect estimation through biologically informed partial pooling, while simultaneously providing modest but consistent improvements in predictive performance.

### 4.2. COPDGene Study Results

We next evaluate the proposed methodology on the COPDGene data. This application illustrates the effect of incorporating curated biochemical subclass information into metabolite effect estimation in a large human metabolomics study.

To illustrate the behavior of the proposed hierarchical estimator, we focus on the regression coefficients corresponding to the *Age, BMI* and *COPD Status* covariates. Similar patterns were observed for the remaining covariates included in the analysis.

Figure 4 summarizes the results for three representative covariates: Age, BMI, and COPD status. Across all three covariates, the Bayesian hierarchical estimator consistently achieved larger RMR values than one, indicating more stable metabolite effect estimates than `MatrixLM`. The greatest improvements occurred at small training sample sizes, where hierarchical borrowing substantially reduced estimation variability. For example, when only 40–60 training subjects were available, the Bayesian estimator produced approximately three to four-fold lower disagreement with the independent reference estimates. As the training sample size increased, the advantage gradually diminished and the RMR approached one, reflecting the increasing precision of the un-pooled `MatrixLM` estimates as more observations became available.

**Fig. 4.**
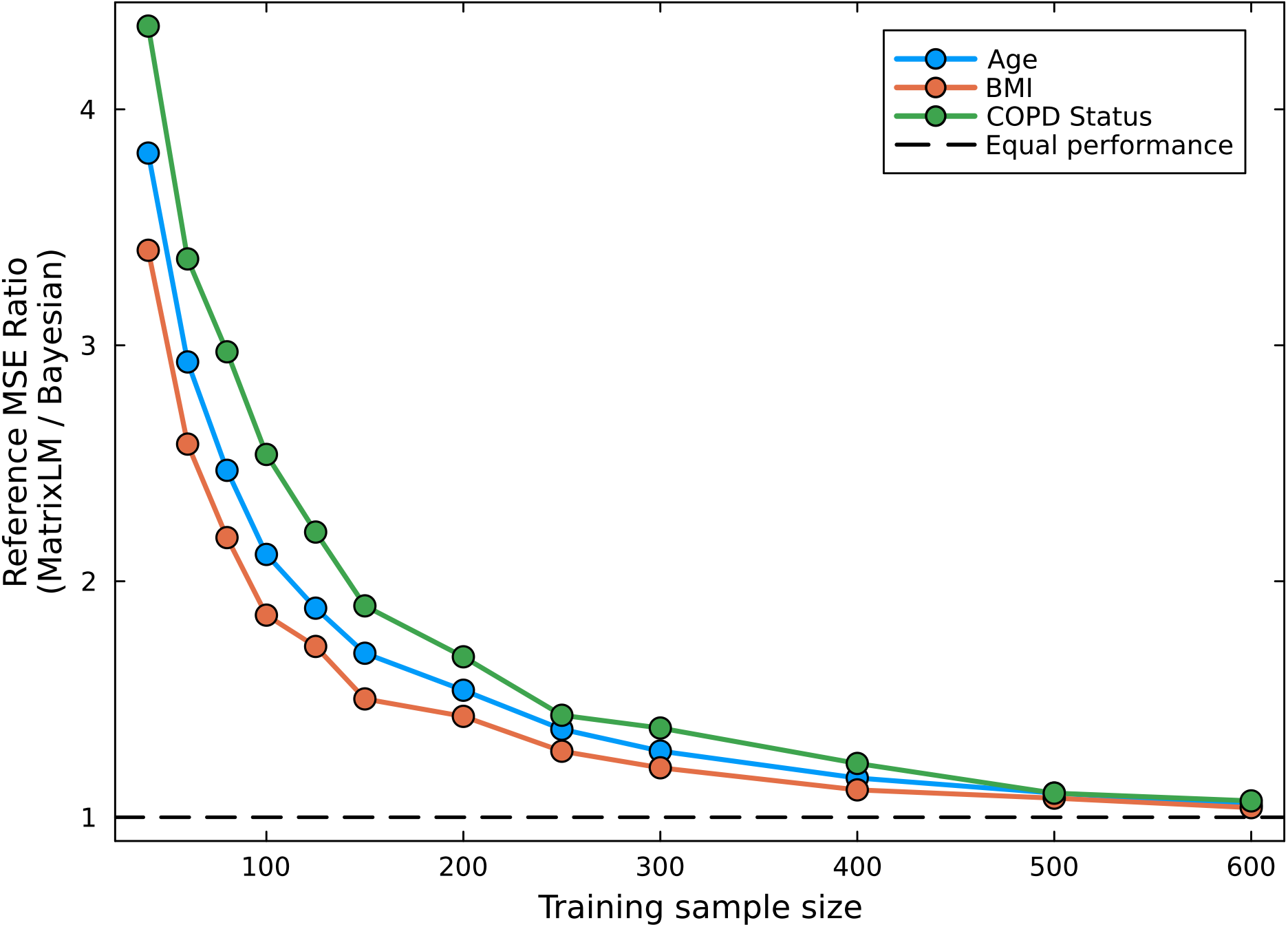
Reference MSE Ratio (RMR) as a function of training sample size for three representative covariates (Age, BMI, and COPD Status) in the COPDGene study. Across all three covariates, the proposed Bayesian hierarchical model consistently achieves higher RMR values than `MatrixLM`, indicating more stable metabolite effect estimates. The greatest improvement is observed at smaller training sample sizes, while the advantage gradually decreases as additional observations become available, reflecting the diminishing need for hierarchical borrowing as estimation precision improves.

These findings closely mirror the behavior observed in the simulation study. The proposed Bayesian framework provides the greatest improvement when estimation is performed using limited data, where borrowing information across biologically related metabolite subclasses effectively reduces sampling variability. As sample size increases, the `MatrixLM` estimates become increasingly stable on their own, naturally reducing the relative benefit of hierarchical partial pooling.

### 4.3. SAMS Study Results

The second application considers the SAMS data, where biologically meaningful triglyceride groupings defined by the number of carbon double bonds and provided the external information used by the hierarchical model. Because the SAMS cohort contains only 98 subjects, we summarize the repeated subsampling analysis in Table 1 rather than using the MSE ratio visualization presented for the simulation study and the COPDGene analysis. Results are shown for the *Interaction SAMS-Fish Oil* covariate, with similar qualitative behavior observed for the remaining covariates.

**Table 1.** Repeated subsampling analysis for the Interaction SAMS-Fish Oil covariate in the SAMS study dataset. For each training sample size, repeated random training subsets were used to estimate metabolite effects using `MatrixLM` and the proposed Bayesian hierarchical model. Agreement with independent reference estimates was summarized using the Reference MSE Ratio (RMR). Across all training sample sizes, the Bayesian hierarchical model consistently produced more stable metabolite effect estimates than `MatrixLM`, with the greatest improvements observed when fewer training samples were available.

| Training Size | Reference Size | MatrixLM MSE | Bayesian MSE | Reference MSE Ratio | Error Reduction (%) |
| --- | --- | --- | --- | --- | --- |
| 20 | 78 | 0.0757 | 0.0309 | 2.4490 | 59.2 |
| 30 | 68 | 0.0534 | 0.0258 | 2.0691 | 51.7 |
| 40 | 58 | 0.0465 | 0.0253 | 1.8401 | 45.7 |
| 50 | 48 | 0.0458 | 0.0295 | 1.5539 | 35.6 |
| 60 | 38 | 0.0477 | 0.0328 | 1.4517 | 31.1 |

Table 1 demonstrates that the proposed Bayesian hierarchical model consistently improved metabolite effect estimation across all training sample sizes. The Reference MSE Ratio exceeded one in every subsampling scenario, indicating that the Bayesian posterior mean estimates agreed more closely with the independent reference estimates than those obtained directly from `MatrixLM`. The largest improvements occurred for the smallest training subsets, where the Bayesian hierarchical model achieved a Reference MSE Ratio of 2.449 and an accompanying Error Reduction of approximately 60%. The latter represents the percentage decrease in mean squared error relative to the independent reference estimates compared with `MatrixLM`.

As the training sample size increased, the Reference MSE Ratio gradually approached one and the corresponding reduction in estimation error became smaller. This behavior closely mirrors the trends observed in the simulation study and the COPDGene analysis. When fewer observations are available, metabolite-specific estimates are more variable and benefit substantially from borrowing information across biologically related triglycerides. As additional observations become available, the `MatrixLM` estimates become increasingly stable, naturally reducing the relative advantage of hierarchical shrinkage.

These results demonstrate that the proposed framework remains effective even when only coarse biological annotations are available. Despite grouping metabolites using only four triglyceride classes defined by degree of unsaturation, biologically informed partial pooling consistently produced more stable metabolite effect estimates across all subsampling scenarios.

### 4.4. PANSteatitis Study Results

Finally, we consider the PANSteatitis study, which demonstrates the proposed methodology in an environmental lipidomics application using curated biochemical subclass annotations. Because the PANSteatitis cohort contains only 44 subjects, we summarize the repeated subsampling analysis in Table 2 rather than using the MSE ratio visualization presented for the simulation study and the COPDGene analysis. Results are shown for the *Weight* covariate, with similar qualitative behavior observed for the remaining covariates.

**Table 2.** Repeated subsampling analysis for the Weight covariate in the PANSteatitis study. For each training sample size, repeated random training subsets were used to estimate metabolite effects using `MatrixLM` and the proposed Bayesian hierarchical model, and agreement with independent reference estimates was summarized using the Reference MSE Ratio (RMR). Across all training sample sizes, the Bayesian hierarchical model consistently produced more stable metabolite effect estimates than `MatrixLM`, with the greatest improvements observed when fewer training samples were available.

| Training Size | Reference Size | MatrixLM MSE | Bayesian MSE | Reference MSE Ratio | Error Reduction (%) |
| --- | --- | --- | --- | --- | --- |
| 10 | 34 | 5.5221 | 2.7468 | 2.0103 | 50.3 |
| 15 | 29 | 2.5236 | 1.0249 | 2.4624 | 59.4 |
| 20 | 24 | 1.8240 | 0.9268 | 1.9681 | 49.2 |
| 25 | 19 | 2.1096 | 1.2774 | 1.6514 | 39.4 |

Table 2 demonstrates that the proposed Bayesian hierarchical model consistently improved metabolite effect estimation across all training sample sizes. The Reference MSE Ratio exceeded one in every subsampling scenario, indicating that the Bayesian posterior mean estimates agreed more closely with the independent reference estimates than those obtained directly from `MatrixLM`. The largest improvements occurred for the smallest training subsets, where the Bayesian hierarchical model achieved a Reference MSE Ratio of 2.462 and an accompanying Error Reduction of approximately 60%.

As in the simulation study, the relative advantage of the Bayesian estimator decreased as the training sample size increased, reflecting the increasing precision of metabolite-specific estimates obtained directly from the observed data. Nevertheless, the Bayesian estimator consistently outper-formed `MatrixLM` across every subsampling scenario, despite the limited sample size available for model fitting. The PANSteatitis application represents the most challenging real-data setting considered in this study, both in terms of sample size and the amount of information available for estimating metabolite effects. The consistent improvement observed across all subsampling scenarios demonstrates that biologically informed partial pooling remains effective even in very small metabolomics studies. Together with the COPDGene and SAMS analyses, these results indicate that the proposed Bayesian framework provides stable improvements in metabolite effect estimation across studies differing substantially in sample size, metabolite composition, and the availability of external biological annotations.

## 5. Discussion

The principal motivation of this work was to demonstrate that external biological information can be incorporated directly into statistical estimation to improve metabolite effect estimation and downstream prediction. Existing approaches estimate regression coefficients for each metabolite independently, even though many metabolites participate in common biochemical pathways or belong to closely related chemical subclasses. Using this biological information can help us get more accurate estimates, especially if the sample size is low and metabolites within a group are tightly connected. The Bayesian hierarchical framework proposed in this paper addresses this limitation by introducing subclass-specific latent effects through which biologically related metabolites borrow information during estimation. Importantly, metabolite-specific effects remain conditionally independent given these latent subclass parameters, preserving interpretable metabolite-level inference while reducing estimation variability.

The simulation study demonstrated that incorporating external biological information through the proposed Bayesian hierarchical framework consistently improved metabolite effect estimation across a broad range of sample sizes and hierarchical heterogeneity regimes. The greatest improvements were observed when sample sizes were small, where biologically informed partial pooling substantially reduced estimation error relative to `MatrixLM`. As the sample size increased, the advantage of the hierarchical model gradually diminished, reflecting the increasing precision of metabolite-specific estimates obtained directly from the observed data. This behavior is consistent with the theoretical motivation for hierarchical modeling; partial pooling provides the greatest benefit when individual estimates are noisy but biologically related.

The three metabolomics applications demonstrated that these improvements extend beyond controlled simulation settings. Across repeated subsampling analyses, the proposed Bayesian framework consistently produced more stable metabolite effect estimates than `MatrixLM`, with the largest improvements observed for the smallest training subsets. As in the simulation study, the relative advantage of hierarchical borrowing decreased as additional observations became available, illustrating that the proposed framework adapts naturally to the amount of information contained in the data. Although improvements in predictive performance were generally more modest, they were consistently observed across the simulation study and real-data applications, indicating that improved metabolite effect estimation can also translate into more reliable prediction.

An important finding is that the proposed framework remained effective across metabolomics studies with markedly different biological annotation structures (Table 3). The COPDGene study utilized 106 metabolite subclasses, the PANSteatitis study contained 13 curated biochemical sub-classes, whereas the SAMS study relied on only four biologically defined triglyceride groups based on degree of unsaturation. Despite these substantial differences in annotation granularity, biologically informed partial pooling consistently improved metabolite effect estimation. These findings suggest that the proposed framework is not restricted to highly detailed biochemical annotations, but can effectively leverage external biological information across multiple levels of biological resolution.

**Table 3.** Characteristics of the metabolomics datasets used to evaluate the proposed Bayesian hierarchical model.

| Dataset | Individuals | Metabolites | Biological Groups | Grouping Variable |
| --- | --- | --- | --- | --- |
| COPDGene | 784 | 999 | 106 | Biochemical subclasses |
| PANSteatitis | 44 | 590 | 13 | Biochemical subclasses |
| SAMS | 98 | 770 | 4 | Number of carbon double bonds |

A limitation of the proposed framework is that it assumes the input metabolite effect estimates are unbiased for their underlying metabolite-specific effects and that their estimation errors are approximately uncorrelated across metabolites. In addition, the effectiveness of biologically informed partial pooling depends on the quality of the external biological annotations used to define metabolite groups. Although the framework remained robust across datasets with substantially different annotation granularities, inaccurate or incomplete groupings may reduce the benefit of information sharing. Finally, as with other hierarchical shrinkage methods, the proposed approach introduces a bias-variance trade-off, whereby borrowing information across biologically related metabolites can diminish truly extreme metabolite effect estimates while reducing estimation variability. Extending the framework to accommodate correlated estimation errors, richer representations of biological similarity, and more flexible hierarchical structures represents an important direction for future research.

Although the present paper focuses on a single-level biological hierarchy, the proposed Bayesian framework readily accommodates more complex hierarchical structures. In preliminary investigations, we implemented a two-level hierarchical model that incorporated multiple levels of metabolite annotation. While this richer hierarchy produced modest improvements over the single-level model, the empirical gains were relatively small compared with the additional computational complexity and increased model specification required. Consequently, we elected to present the simpler one-level hierarchy, which provides a favorable balance between interpretability, computational efficiency, and estimation performance. Nevertheless, many metabolomics databases naturally organize metabolites into nested biological ontologies consisting of superclasses, subclasses, and individual metabolites. The proposed framework can therefore be extended in a straightforward manner to accommodate multiple hierarchical levels whenever the scientific application warrants the additional model complexity.

Future work could explore the incorporation of more complex representations of external biological information beyond manually curated metabolite subclasses. Rather than relying on discrete biological groupings, the proposed hierarchical framework could be extended to leverage continuous measures of similarity derived from molecular structure, biological pathways, interaction networks, or learned feature representations. Such representations have the potential to capture more nuanced relationships among metabolites and may further improve the effectiveness of biologically informed partial pooling, particularly in settings where curated annotations are incomplete or unavailable.

Of course, the proposed framework is not restricted to metabolomics. Because it operates as a post-estimation procedure requiring only feature-specific effect estimates and their associated standard errors, it can be applied following a wide range of statistical estimation methods. Consequently, similar hierarchical borrowing strategies could be employed in other high-dimensional biological applications, including proteomics, transcriptomics, microbiome studies, and related multi-omics settings wherever meaningful external feature information is available.

In summary, the proposed Bayesian hierarchical framework provides a principled approach for incorporating external biological information into metabolite effect estimation. Across both simulated and real metabolomics studies, the method consistently produced more stable and reproducible metabolite effect estimates, with the greatest improvements observed when sample sizes were limited. Although gains in predictive performance were more modest, they were consistently observed in both simulated and real data, demonstrating that improved effect estimation can also translate into more reliable prediction. More broadly, the proposed framework illustrates a general strategy for integrating external biological information into statistical estimation. In this work, external information is represented through curated metabolite subclasses, but the framework is equally applicable to richer representations of biological relationships as they become available. We believe that integrating structured biological knowledge with statistical inference provides a flexible and broadly applicable framework for improving estimation in modern high-dimensional omics studies.

## 6. Data and Code Availability

The Pansteatatis Mozambique Tilapia data are available at the NIH Common Fund’s National Metabolomics Data Repository (NMDR), the Metabolomics Workbench (https://www.metabolomicsworkbench.org; accessed on 21 July 2026), where they have been assigned Project ID PR000705. The data can be accessed directly via the Project DOI: 10.21228/M8JH5X. COPDGene and SPIROMICS data are available at the NIH Common Fund’s National Metabolomics Data Repository (NMDR), the Metabolomics Workbench (https://www.metabolomicsworkbench.org; accessed on 21 July 2026), where they have been assigned Project ID PR001048. The data can be accessed directly via the Project DOI: 10.21228/M87D6G.

The code used to generate the figures and tables presented in this study is available at https://github.com/senresearch/BayesHierarchy_MLM. The MatrixLM.jl package is available at https://github.com/senresearch/MatrixLM.jl.

The MetabolomicsWorkbenchAPI.jl package is available at https://github.com/senresearch/MetabolomicsWorkbenchAPI.jl.

## A. Appendix

### A.1. Prior Specification

This section describes the prior distributions assigned to the unknown parameters of the Bayesian hierarchical model introduced in Section 2.2. To complete the Bayesian hierarchical model, prior distributions are assigned to the global mean parameter and the variance components governing within-group and between-group heterogeneity.

The global mean effect is assigned a normal prior,

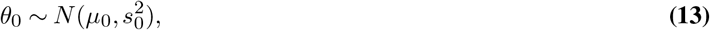

where *µ*_0_ and 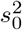 are fixed hyperparameters chosen to represent prior beliefs regarding the overall magnitude of metabolite effects. In the absence of strong prior information, *µ*_0_ may be set to zero and 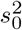 chosen sufficiently large to yield a weakly informative prior.

The variance parameters 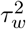 and 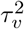 play a central role in determining the degree of shrinkage induced by the hierarchy. Specifically, 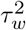 controls the variability of metabolite effects within a biological group, while 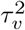 controls the variability of group-level effects around the global mean. Smaller values correspond to stronger pooling, whereas larger values allow greater heterogeneity among effects.

Rather than assigning inverse-gamma priors directly to these variance components, we adopt Half-Cauchy priors on the corresponding standard deviation parameters,

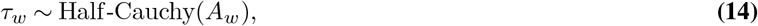

and

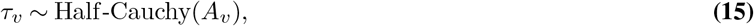

where *A*_*w*_ and *A*_*v*_ are positive scale parameters.

The Half-Cauchy distribution has become a widely used default prior for hierarchical variance components because it combines substantial mass near zero with heavy tails (16, 30). Consequently, the prior permits strong shrinkage when supported by the data while still allowing large variance values when substantial heterogeneity is present. This behavior avoids the excessive influence that can arise from overly informative variance priors and often yields more robust inference in hierarchical models. Following common practice, we use the same scale parameter for both variance components,

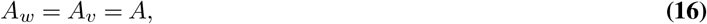

although the framework readily accommodates different choices when prior knowledge suggests distinct levels of within-group and between-group variability.

For posterior computation, the Half-Cauchy priors are represented using an equivalent inverse-gamma scale-mixture formulation. This representation preserves the desired prior distribution while producing conditionally conjugate updates that facilitate efficient Gibbs sampling. Details of the resulting posterior computation is outlined in Appendix A.2.

### A.2. Posterior Inference

This section describes the posterior computation of our proposed Bayesian hierarchical model. Let

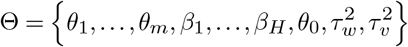

denote the collection of unknown model parameters. Posterior inference is based on the distribution of Θ conditional on the observed MLM effect estimates

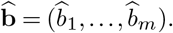

Combining the likelihood in Equation (1) with the hierarchical model specified in Equations (2)–(3) and the prior distributions in Equations (13)–(15), Bayes’ theorem yields the joint posterior distribution

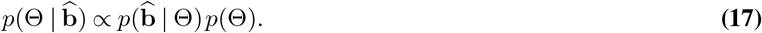

Expanding the individual components gives

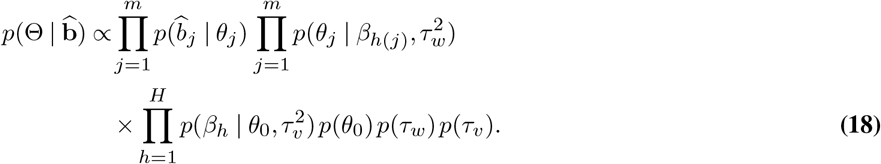

Although each level of the hierarchy involves normal distributions, the Half-Cauchy priors assigned to the variance parameters prevent the posterior distribution from having a closed-form analytical solution. Consequently, direct evaluation of posterior moments and marginal distributions is generally infeasible.

To facilitate computation, we adopt the inverse-gamma scale-mixture representation of the Half-Cauchy distribution. Following (16), a Half-Cauchy prior on a scale parameter may be expressed as a hierarchical mixture involving inverse-gamma random variables. Specifically,

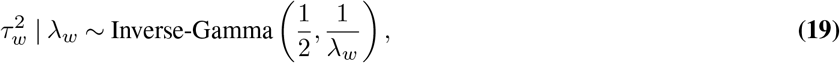

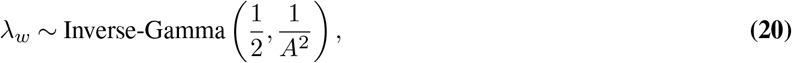

and similarly,

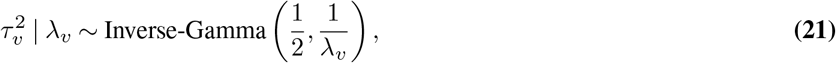

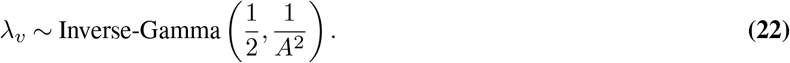

Integrating out the auxiliary variables *λ*_*w*_ and *λ*_*v*_ recovers the original Half-Cauchy priors specified in Appendix A.1. This representation introduces latent variables but yields conditionally conjugate posterior distributions for all unknown quantities in the hierarchy.

The resulting conditional distributions form the basis of an efficient Gibbs sampling algorithm, described in the following subsection. Complete derivations of all full conditional distributions are provided in Appendix A.3.

### A.3. Derivation of Full Conditional Distributions

This section derives the full conditional distributions used in the Gibbs sampling algorithm described in Appendix A.4. The Half-Cauchy priors on scale parameters follow the hierarchical variance-prior recommendations of Gelman (30), and the inverse-gamma auxiliary-variable representation follows Makalic and Schmidt (16). The resulting Gibbs sampler is a standard MCMC procedure for approximating posterior distributions (20).

For notational convenience, let *h*(*j*) denote the biological group membership of metabolite *j*, and let

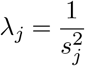

denote the observation-level precision associated with the MatrixLM estimate 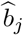.

The Bayesian hierarchical model is

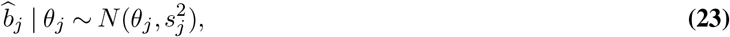

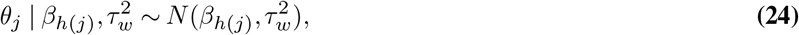

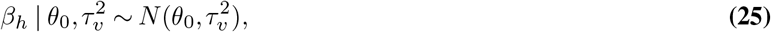

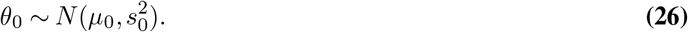

The Half-Cauchy priors on *τ*_*w*_ and *τ*_*v*_ are represented as

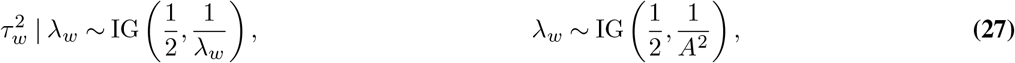

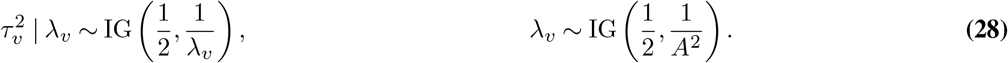

Here, IG(*a, b*) denotes the inverse-gamma distribution with density proportional to

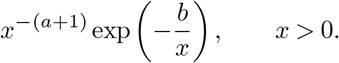

#### A.3.1. Full Conditional for θ_j_

The full conditional distribution of *θ*_*j*_ depends on the MatrixLM estimate 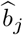 and the group-level parameter *β*_*h*(*j*)_. Up to proportionality,

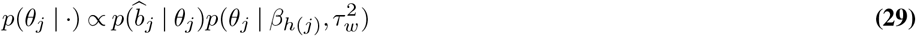

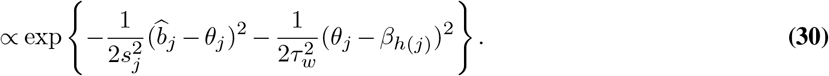

Collecting terms in *θ*_*j*_ gives a normal full conditional distribution,

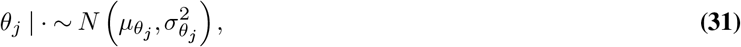

where

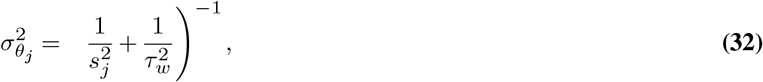

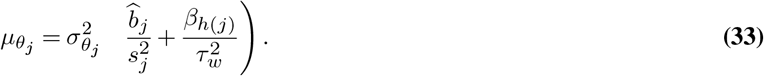

Equivalently, using 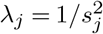,

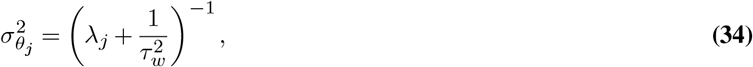

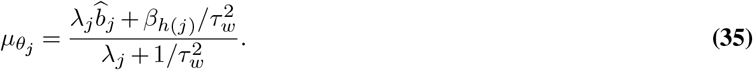

#### A.3.2. Full Conditional for β_h_

Let

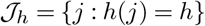

denote the set of metabolites assigned to group *h*, and let *m*_*h*_ = |*J*_*h*_|. The full conditional distribution of *β*_*h*_ depends on the metabolite-level effects in group *h* and on the global mean *θ*_0_:

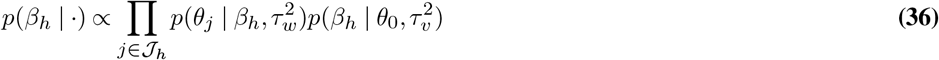

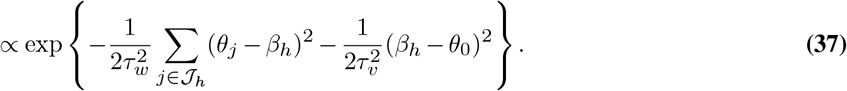

Thus,

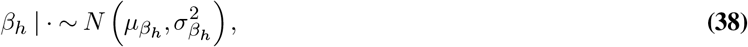

where

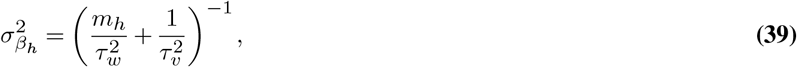

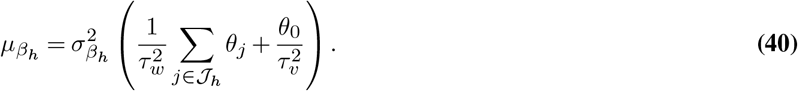

#### A.3.3. Full Conditional for θ_0_

The full conditional distribution of *θ*_0_ depends on the group-level effects *β*_1_, …, *β*_*H*_ and the prior distribution for *θ*_0_:

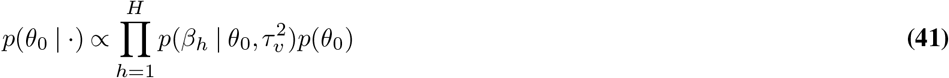

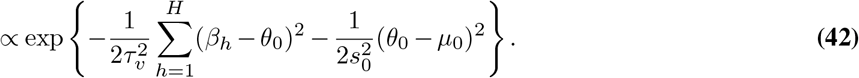

Therefore,

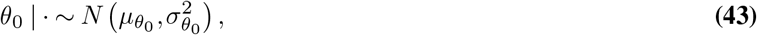

where

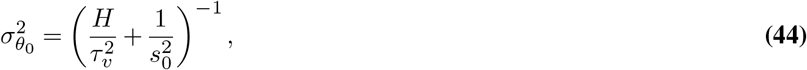

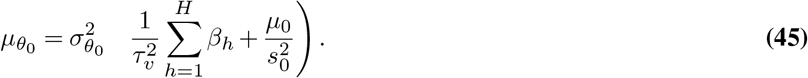

#### A.3.4. Full Conditional for 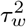

The full conditional distribution of 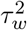 depends on the deviations of the metabolite-level effects from their corresponding group means and on the auxiliary variable *λ*_*w*_.

Define the within-group sum of squares

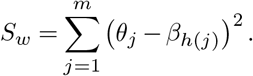

Then,

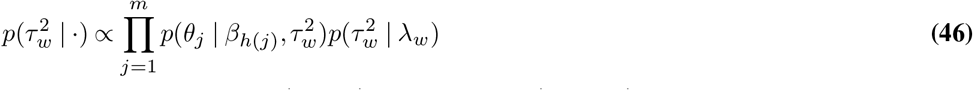

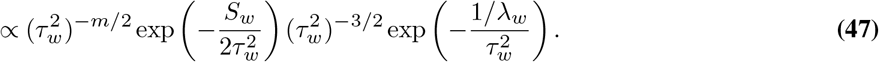

Combining terms gives

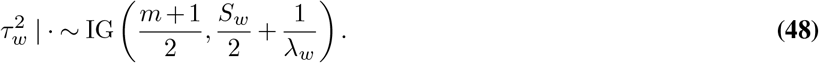

#### A.3.5. Full Conditional for 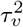

The full conditional distribution of 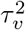 depends on the deviations of the group-level effects from the global mean and on the auxiliary variable *λ*_*v*_.

Define the between-group sum of squares

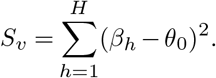

Then,

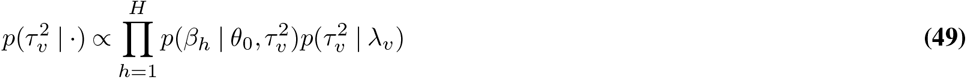

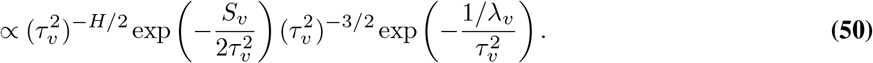

Therefore,

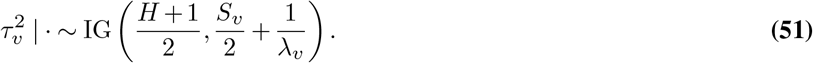

#### A.3.6. Full Conditional for λ_w_

The full conditional distribution of *λ*_*w*_ depends on the conditional prior for 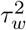 and the prior for *λ*_*w*_:

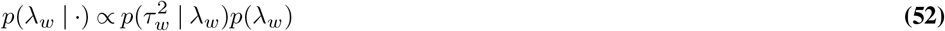

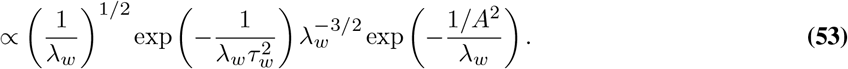

Combining powers of *λ*_*w*_ gives

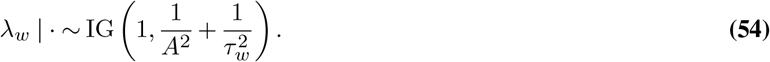

#### A.3.7. Full Conditional for λ_v_

Analogously, the full conditional distribution of *λ*_*v*_ is

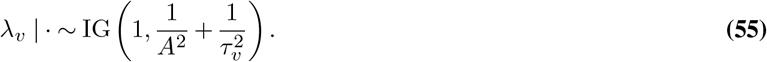

### A.4. Gibbs Sampling Algorithm

The full conditional distributions derived in Appendix A.3 lead to a straightforward Gibbs sampling algorithm for posterior computation.

The inverse-gamma scale-mixture representation introduced in Appendix A.2 yields conditionally conjugate posterior distributions for all unknown parameters, allowing each parameter to be sampled directly from its full conditional distribution.

Let

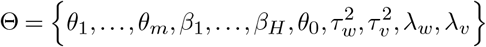

denote the complete set of unknown quantities after introducing the auxiliary scale-mixture variables. The Gibbs sampler iteratively updates each parameter conditional on the current values of all remaining parameters.

At iteration *t*, the following updates are performed:

1. Update the metabolite-specific effects

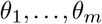

from their full conditional distributions.
2. Update the group-level effects

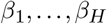

from their full conditional distributions.
3. Update the global mean parameter

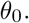
4. Update the within-group variance component

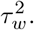
5. Update the between-group variance component

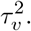
6. Update the auxiliary scale-mixture variables

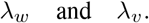

#### Algorithm 1

Gibbs Sampling for the Bayesian Hierarchical Model

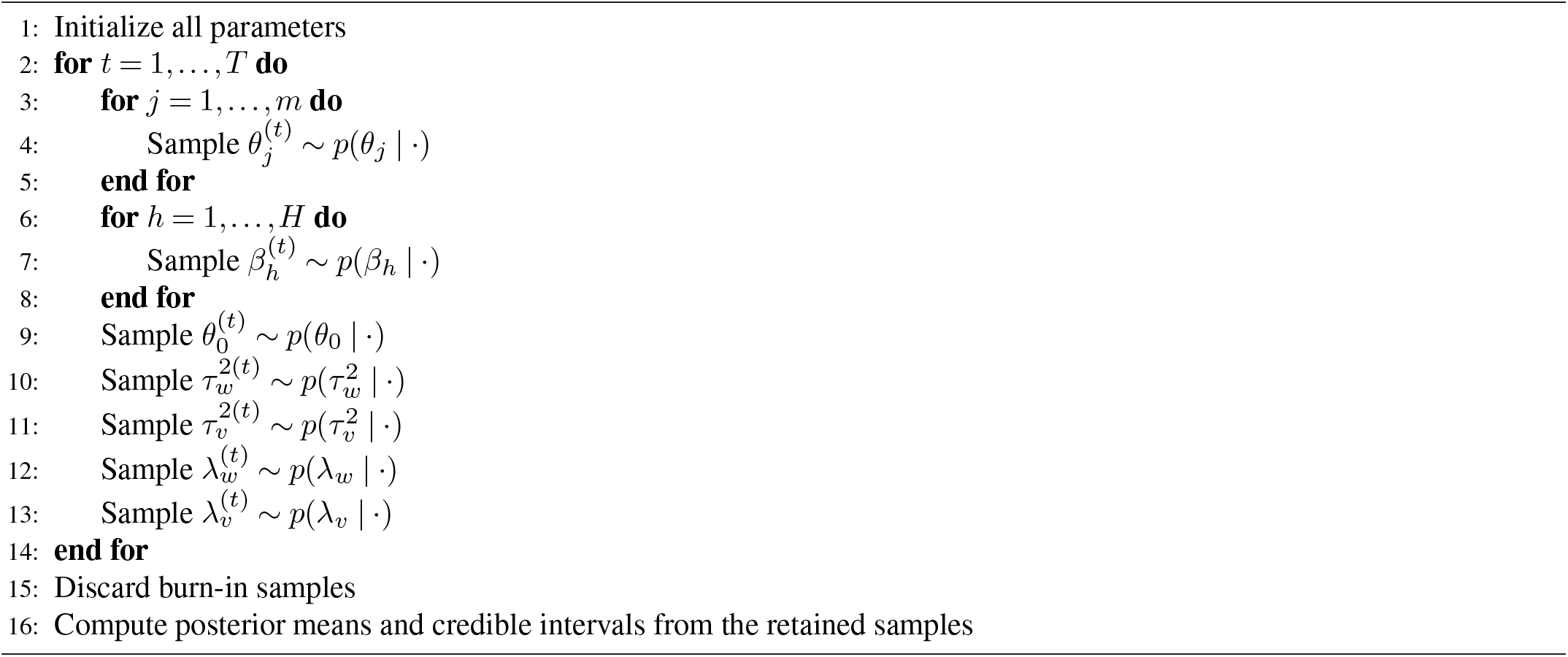

The full conditional distributions for each update are available in closed form and are provided in Appendix A.3. Because all updates are obtained from standard normal or inverse-gamma distributions, posterior sampling is computationally efficient and avoids the need for rejection sampling, Metropolis–Hastings steps, or numerical optimization.

The Gibbs sampler was initialized using the MLM estimates and run for *T* iterations. The first *B* iterations were discarded as burn-in, and posterior summaries were computed using the remaining samples. Convergence was assessed using trace plots and standard Markov chain Monte Carlo diagnostics.

For a generic iteration, the Gibbs sampling procedure is summarized in Algorithm 1.

The resulting posterior samples provide estimates of metabolite-specific effects, group-level effects, variance components, and associated credible intervals.

## Bibliography

1. Oliver Fiehn. Metabolomics–the link between genotypes and phenotypes. Plant molecular biology, 48(1):155–171, 2002.

2. Gary J Patti, Oscar Yanes, and Gary Siuzdak. Metabolomics: the apogee of the omic triology. Nature reviews. Molecular cell biology, 13(4):263, 2012.

3. Warwick B Dunn, David Broadhurst, Paul Begley, Eva Zelena, Sue Francis-McIntyre, Nadine Anderson, Marie Brown, Joshau D Knowles, Antony Halsall, John N Haselden, et al. Procedures for large-scale metabolic profiling of serum and plasma using gas chromatography and liquid chromatography coupled to mass spectrometry. Nature protocols, 6(7): 1060–1083, 2011.

4. Joseph Antonelli, Brian Claggett, Mir Henglin, Jeramie D Watrous, Kim A Lehmann, Pavel Hushcha, Olga Demler, Samia Mora, Teemu Niiranen, Alexandre C Pereira, et al. Statistical methods and workflow for analyzing human metabolomics data. arXiv preprint arXiv:1710.03436, 2017.

5. Najeha R Anwardeen, Ilhame Diboun, Younes Mokrab, Asma A Althani, and Mohamed A Elrayess. Statistical methods and resources for biomarker discovery using metabolomics. BMC bioinformatics, 24(1):250, 2023.

6. Dennis V Lindley and Adrian FM Smith. Bayes estimates for the linear model. Journal of the Royal Statistical Society Series B: Statistical Methodology, 34(1):1–18, 1972.

7. Willard James and Charles Stein. Estimation with quadratic loss. In Jerzy Neyman, editor, Proceedings of the Fourth Berkeley Symposium on Mathematical Statistics and Probability, volume 1, pages 361–379. University of California Press, Berkeley, CA, 1961.

8. Bradley Efron and Carl Morris. Stein’s estimation rule and its competitors—an empirical bayes approach. Journal of the American Statistical Association, 68(341):117–130, 1973.

9. Andrew Gelman and Jennifer Hill. Data analysis using regression and multilevel/hierarchical models. Cambridge university press, 2007.

10. Bradley Efron and Carl Morris. Data analysis using stein’s estimator and its generalizations. Journal of the American Statistical Association, 70(350):311–319, 1975.

11. Bradley P Carlin and Thomas A Louis. Bayesian methods for data analysis. CRC press, 2008.

12. Christopher E. Gillies, Theodore S. Jennaro, Michael A. Puskarich, Ruchi Sharma, Kevin R. Ward, Xudong Fan, Alan E. Jones, and Kathleen A. Stringer. A multilevel bayesian approach to improve effect size estimation in regression modeling of metabolomics data utilizing imputation with uncertainty. Metabolites, 10(8), 2020. ISSN 2218-1989. doi: 10.3390/metabo10080319.

13. Claudio Busatto and Mark A Van De Wiel. Informative co-data learning for high-dimensional horseshoe regression. Biometrical Journal, 68(1):e70105, 2026.

14. Marco Molinari, Andrea Cremaschi, Maria De Iorio, Nishi Chaturvedi, Alun Hughes, and Therese Tillin. Bayesian dynamic network modelling: an application to metabolic associations in cardiovascular diseases. Journal of Applied Statistics, 51(1):114–138, 2024.

15. Willem Collier, Benjamin Haaland, Lesley A Inker, Hiddo JL Heerspink, and Tom Greene. Comparing bayesian hierarchical meta-regression methods and evaluating the influence of priors for evaluations of surrogate endpoints on heterogeneous collections of clinical trials. BMC Medical Research Methodology, 24(1):39, 2024.

16. Enes Makalic and Daniel F Schmidt. A simple sampler for the horseshoe estimator. IEEE Signal Processing Letters, 23(1):179–182, 2015.

17. David S Wishart, AnChi Guo, Eponine Oler, Fei Wang, Afia Anjum, Harrison Peters, Raynard Dizon, Zinat Sayeeda, Siyang Tian, Brian L Lee, et al. Hmdb 5.0: the human metabolome database for 2022. Nucleic acids research, 50(D1):D622–D631, 2022.

18. Minoru Kanehisa, Miho Furumichi, Yoko Sato, Mari Ishiguro-Watanabe, and Mao Tanabe. Kegg: integrating viruses and cellular organisms. Nucleic acids research, 49(D1):D545–D551, 2021.

19. Andrew Gelman, John B. Carlin, Hal S. Stern, David B. Dunson, Aki Vehtari, and Donald B. Rubin. Bayesian data analysis. CRC press, 2013.

20. Christian P Robert and George Casella. Monte Carlo statistical methods, volume 2. Springer, 2004.

21. Gregory Farage, Chenhao Zhao, Hyo Young Choi, Timothy J. Garrett, Marshall B. Elam, Katerina Kechris, and Sàunak Sen. Matrix linear models for connecting metabolite composition to individual characteristics. Metabolites, 15(2), 2025. ISSN 2218-1989. doi: 10.3390/metabo15020140.

22. Lucas A. Gillenwater, Katerina J. Kechris, Katherine A. Pratte, Nichole Reisdorph, Irina Petrache, Wassim W. Labaki, Wanda O’Neal, Jerry A. Krishnan, Victor E. Ortega, Dawn L. DeMeo, and Russell P. Bowler. Metabolomic Profiling Reveals Sex Specific Associations with Chronic Obstructive Pulmonary Disease and Emphysema. Metabolites, 11(3):161, mar 2021. ISSN 2218-1989. doi: 10.3390/metabo11030161.

23. Frank Dieterle, Alfred Ross, Götz Schlotterbeck, and Hans Senn. Probabilistic Quotient Normalization as Robust Method to Account for Dilution of Complex Biological Mixtures. Application in 1 H NMR Metabonomics. Analytical Chemistry, 78(13):4281–4290, jul 2006. ISSN 0003-2700, 1520-6882. doi: 10.1021/ac051632c.

24. Tim J Garrett, Michelle A Puchowicz, Qingming Dong, Gregory Farage, Richard Childress, Edwards A Park, Joy Guingab, Claire L Simpson, Saunak Sen, Elizabeth C Brogdon, Logan M Buchanan, Rajendra Raghow, and Marshall B Elam. Effect of Treatment on Metabolites, Lipids and Prostanoids in Patients with Statin Associated Muscle Symptoms (SAMS). PlosOne (in press), December 2023.

25. Cosmin Lazar, Thomas Burger, and Samuel Wieczorek. imputelcmd: a collection of methods for left-censored missing data imputation. R package, version, 2, 2015.

26. Jeffrey T Leek, W Evan Johnson, Hilary S Parker, Andrew E Jaffe, and John D Storey. The sva package for removing batch effects and other unwanted variation in high-throughput experiments. Bioinformatics, 28(6):882–883, 2012.

27. W Evan Johnson, Cheng Li, and Ariel Rabinovic. Adjusting batch effects in microarray expression data using empirical bayes methods. Biostatistics, 8(1):118–127, 2007.

28. Jeremy P Koelmel, Candice Z Ulmer, Susan Fogelson, Christina M Jones, Hannes Botha, Jacqueline T Bangma, Theresa C Guillette, Wilmien J Luus-Powell, Joseph R Sara, Willem J Smit, et al. Lipidomics for wildlife disease etiology and biomarker discovery: a case study of pansteatitis outbreak in south africa. Metabolomics, 15(3):38, 2019.

29. Manish Sud, Eoin Fahy, Dawn Cotter, Kenan Azam, Ilango Vadivelu, Charles Burant, Arthur Edison, Oliver Fiehn, Richard Higashi, K Sreekumaran Nair, et al. Metabolomics workbench: An international repository for metabolomics data and metadata, metabolite standards, protocols, tutorials and training, and analysis tools. Nucleic acids research, 44(D1): D463–D470, 2016.

30. Andrew Gelman. Prior distributions for variance parameters in hierarchical models. Bayesian Analysis, 1(3):515–533, 2006.

